# β2-integrins as biomarkers in urothelial cancer

**DOI:** 10.1101/2025.05.25.656033

**Authors:** Marc Llort Asens, Imran Khan, Susanna Carola Fagerholm

## Abstract

β2-integrins are a family of adhesion proteins expressed in immune cells that play multiple roles in anti-tumor immunity. β2-integrins regulate tumor infiltration of anti-tumorigenic immune cells such as cytotoxic CD8+ T cells and NK cells. However, they also regulate the activity of myeloid cells, such as macrophages, which can have both anti- and pro-tumorigenic properties. The role of β2-integrins in urothelial cancer remains poorly understood. Here, we have investigated the role of different β2-integrins, and their cytoplasmic regulators, in urothelial cancer, by utilizing RNA expression data. We found that ITGAL (encoding for CD11a) and FERMT3 (encoding for the integrin regulator kindlin-3) have a positive correlation with patient survival. EcoTyper analysis revealed increased infiltration of CD8+ T cells and NK cells in ITGAL high samples, but ITGAL or FERMT3 expression did not correlate with response to immunotherapy. In contrast, ITGAM and ITGAX (which encode for myeloid markers CD11b and CD11c) and FLNA (encoding for the integrin regulator filamin A) correlated with poor survival and reduced responsiveness to immunotherapy and critically regulate the tumor myeloid immune landscape (M1/M2 macrophages, cDC1 dendritic cells). Therefore, different β2-integrins may be used as biomarkers to differentiate urothelial cancer patients with different immune landscapes, responding differently to therapy.

## Introduction

Cancer remains a leading global health challenge, nearly 20 million people worldwide were diagnosed with cancer in 2020. Urothelial bladder cancer was ranked 9^th^ in incidence and 13^th^ in cancer related mortality as per World Health Organization, accounting to over 614,000 cases and more than 220,000 deaths (1) As for other cancer types, immunotherapy is used also in urothelial bladder cancer, but response rates lie below 30% (2,3). This highlights an urgent need for better understanding of molecular mechanisms behind urothelial bladder cancer development, characterization of the immune landscape of the disease, and identification of reliable novel biomarkers predicting disease progression and response to therapy.

β2-integrins are a family of transmembrane heterodimeric proteins located at the surface of all leukocytes (4). They play an essential role in immune cell trafficking to inflammation sites and tumors by allowing leukocytes to move between the blood stream and the surrounding tissues (5). Among many other functions, they are also key to the modulation of the immune system, by regulating the activation state and polarization of myeloid cells such as macrophages and dendritic cells. β2-integrins are also central in the formation of immunological synapses, complex structures that immune cells form to communicate between themselves and to mediate tumor cell killing (5).

Despite sharing a common β subunit, encoded by the gene ITGB2, β2-integrins differ in their α subunits and their expression can be restricted to different cell types (4). Thus, the α subunit encoded by the gene ITGAL pairs with β2 to form the integrin αLβ2 (also known as LFA-1), which is expressed in all leukocytes and is the predominant β2-integrin in lymphocytes. The α subunit encoded by the gene ITGAM forms the integrin αMβ2 (also called Mac-1) and is mostly expressed in myeloid cells (particularly in neutrophils, but also in macrophages and dendritic cells). The α subunit encoded by the gene ITGAX forms the integrin αXβ2 (CD11c/CD18), which is most abundant in dendritic cells. Lastly, the α subunit encoded by the gene ITGAD encodes for the integrin αDβ2, the most recently discovered and least known β2-integrin, which is expressed in neutrophils, monocytes and natural killer cells.

Integrins are regulated by interactions with cytoplasmic proteins binding to their cytoplasmic domains (tails). For example, talin (encoded by TLN1) and kindlin-3 (encoded by FERMT3) are essential for integrin activation and binding to ligands (6,7)(8–10), whilst filamin A (encoded by FLNA) has been reported to play both positive and negative roles in integrin function (11)(12)

Due to their heterogeneity in both structure and expression profile, it is not surprising that different β2-integrins perform different functions, at times even completely opposite roles in the modulation of the immune system both in health and in disease. Hence, while αLβ2 is key to trafficking of cytotoxic CD8+ T cells into tumors and CD8+ T cell effector functions(5), αMβ2 can play an opposite role in myeloid cells such as macrophages and dendritic cells (DCs) by suppressing their activation, function as antigen presenting cells, activators of T cells and anti-tumor responses (13–16). However, the role of β2-integrins in regulating urothelial bladder cancer immune landscape and/or responsiveness to therapy remains poorly understood.

Here, we have investigated the role of β2-integrins and their cytoplasmic regulators in urothelial cancer, by utilizing the IMvigor210 cohort (NCT02108652). We found that although expression of ITGAL and FERMT3 (which regulate immune cell infiltration into tumors) do correlate with increased survival, they do not predict response to immunotherapy. In contrast, myeloid markers ITGAM and ITGAX were inversely correlated with patient survival and response to immunotherapy, and also critically regulate the myeloid immune landscape of the tumors. Therefore, different subclasses of β2-integrins may be used as biomarkers to predict disease outcome and responsiveness to immunotherapy in urothelial cancer.

## Results

### T cell markers expression levels effect on survival and response to immunotherapy

We investigated whether expression levels of ITGB2, ITGAL, and FERMT3, which are important for leukocyte (T cell, NK cell) infiltration into tumors, had an effect on cancer patient survival. To that aim, we used the publicly available gene expression data from the IMvigor210 study, which evaluates the effect of anti-PD-L1 immunotherapy on survival of patients of urothelial cancer. We categorized the patients into two separate groups (high or low) based on individual gene expression levels of ITGB2, ITGAL, and FERMT3 (see Materials and Methods). We observed no significant differences in the length of overall survival between patients with high or low expression of ITGB2, which encodes for the common β2-chain of all four β2-integrins (Fig.1A). In contrast, we identified that patients with a high expression of ITGAL (ITGAL high group) showed a significantly longer overall survival than the patients with low ITGAL expression (ITGAL low) [P=0.013] and that their group reached the median of survival probability remarkably later (5 months) than the group with low ITGAL expression (Fig. 1B). In line with these findings, we also identified that patients with a high expression of FERMT3 (FERMT3 high) showed a significantly longer overall survival than the patients with low FERMT3 expression (FERMT3 low) [P=0.043] (Fig. 1C).

**Fig. 1.**
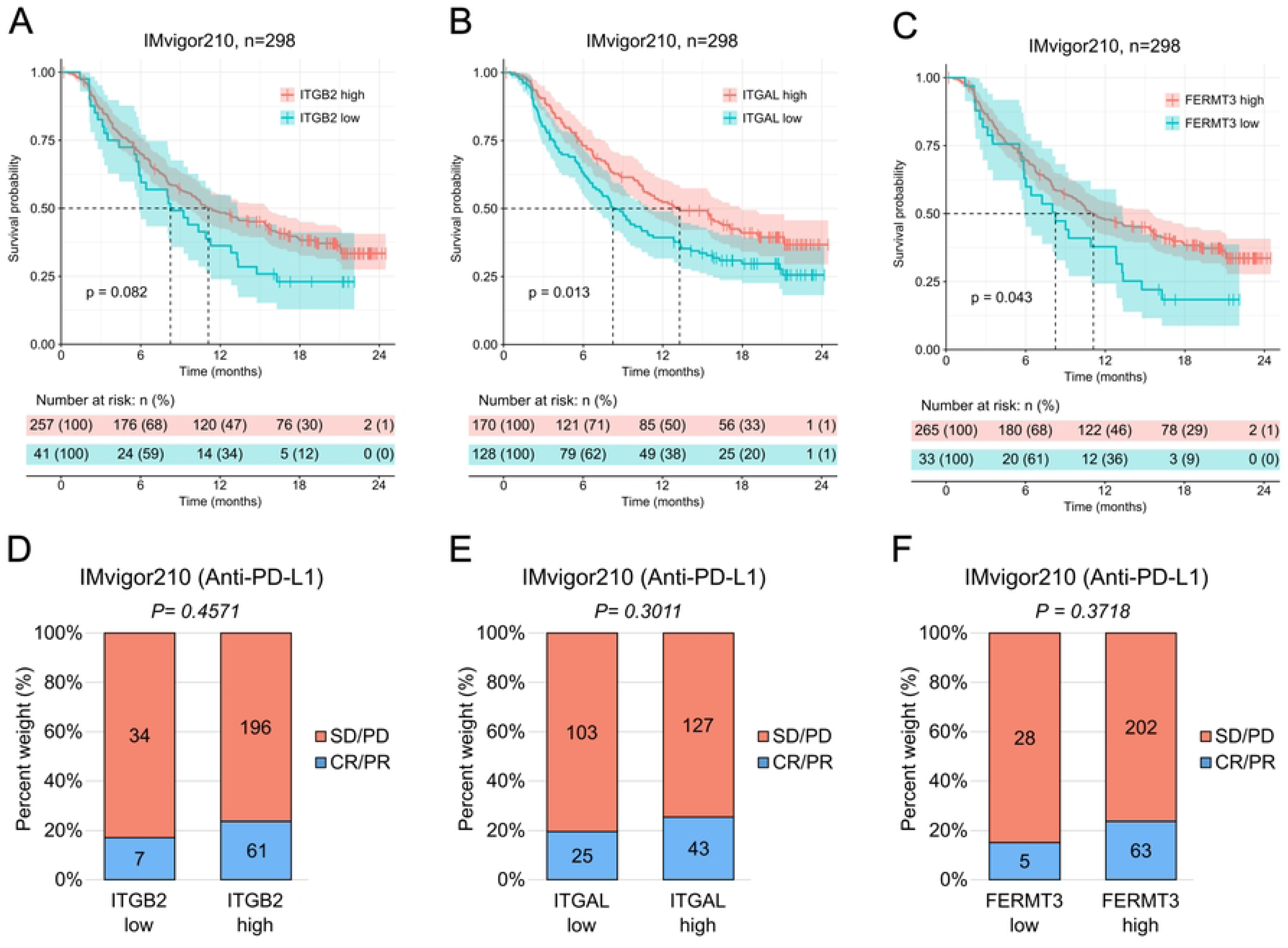
High expression levels of ITGAL and FERMT3 correlate with longer overall survival in urothelial cancer patients. **A-C**, effect of gene expression in infiltrating leukocytes. Kaplan-Meier estimator of the overall survival after treatment with anti-PD-L1 in the IMvigor210 cohort of ITGB2 high (n=257) and ITGB2 low (n=41) patients (**A**), ITGAL high (n=170) and ITGAL low (n=128) patients (**B**), and FERMT3 high (n=265) and FERMT3 low (n=33) patients (**C**). The dotted lines indicate the time at which each group reached median survival (ITGB2 high: 11.1 months, ITGB2 low: 8.25 months; ITGAL high: 13.27 months, ITGAL low: 8.25 months; FERMT3 high: 11.1 months, FERMT3 low: 8.25 months)

However, the expression levels of ITGB2, ITGAL or FERMT3 did not significantly affect how well the patients responded to anti-PD-L1 immunotherapy as accounted by the number of patients with stable disease or progressive disease (SD/PD) versus the number of patients with a complete response or partial response (CR/PR) (Fig. 1D-F). Together, these results highlight the importance of ITGAL and FERMT3 in the immune response against urothelial cancer.

The colored vertical marks on the plot indicate censored events for each group in time. Below the plot, risk table indicating the number of patients at risk belonging to each group at each point in time: high, in orange, or low, in blue. Statistical analysis was done using the log-rank (Mantel-Cox) test. P-values are shown on the plots. **D-F**, rate of clinical response to anti-PD-L1 immunotherapy in the IMvigor210 cohort (SD/PD, stable disease/progressive disease, n=257 (**D**), n=170 (**E**), n=265 (**F**); CR/PR, complete response/partial response, n=41 (**D**), n=128 (**E**), n=33 (**F**)). Statistical analysis was done using the chi-squared test. P-values are shown on the plots.

### Myeloid **β**2-integrin expression levels impact on survival and immunotherapy response

Next, we wanted to study whether the expression levels of ITGAM, ITGAX and ITGAD, which are typically expressed in myeloid cells and are associated with suppression of the immune response, affected patient survival and response to anti-PD-L1 immunotherapy in the urothelial cancer cohort.

We found that patients with low levels of ITGAM (ITGAM low) or ITGAX (ITGAX low), analyzed independently, showed significantly longer overall survival compared to those with high levels of expression (ITGAM high, and ITGAX high, respectively) (Fig. 2A-B). Additionally, they responded significantly better to anti-PD-L1 immunotherapy (Fig. 2D-E).

**Fig. 2.**
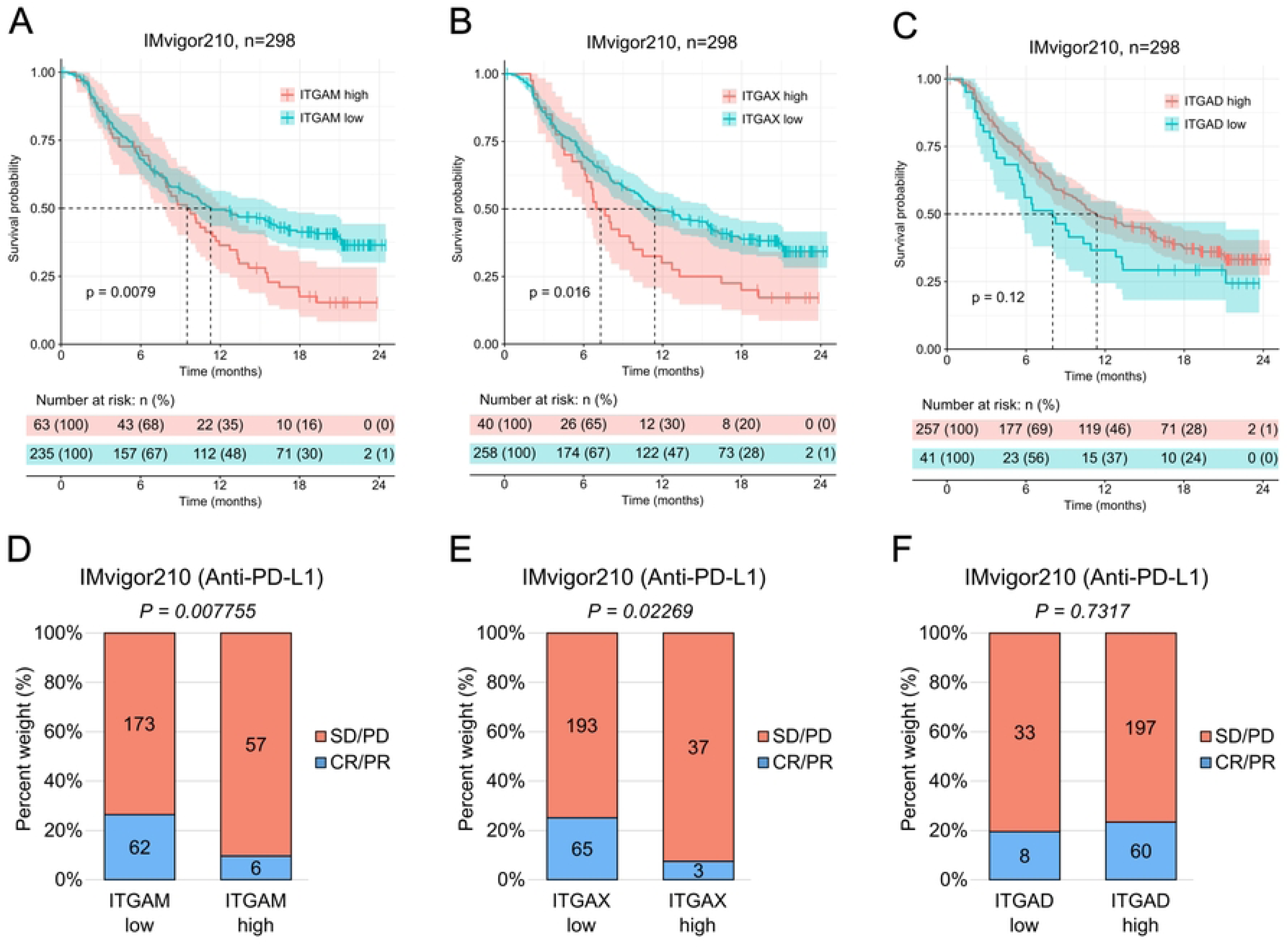
Low expression levels of ITGAM and ITGAX correlate with longer overall survival in urothelial cancer patients and enhanced response to anti-PD-L1 immunotherapy. **A-C**, Kaplan-Meier estimator of the overall survival after treatment with anti-PD-L1 of ITGAM high (n=63) and ITGAM low (n=235 patients (**A**), ITGAX high (n=40) and ITGAX low (n=258) patients (**B**), and ITGAD high (n=257) and ITGAD low (n=41) patients (**C**). The dotted lines indicate the time at which each group reached median survival (ITGAM high: 9.49 months, ITGAM low: 11.27 months; ITGAX high: 7.29 months, ITGAX low: 11.4 months; ITGAD high: 11.36 months, ITGAD low: 8.02 months). The colored vertical marks on the plot indicate censored events for each group in time. Below the plot, risk table indicating the number of patients at risk belonging to each group at each point in time: high, in orange, or low, in blue. Statistical analysis was done using the log-rank (Mantel-Cox) test. P-values are shown on the plots. **D-F**, rate of clinical response to anti-PD-L1 immunotherapy (SD/PD, stable disease/progressive disease, n=63 (**D**), n=40 (**E**), n=257 (**F**); CR/PR, complete response/partial response, n=235 (**D**), n=258 (**E**), n=41 (**F**)). Statistical analysis was done using the chi-squared test. P-values are shown on the plots.

In contrast, the expression levels of ITGAD did not seem to alter significantly either the overall survival (Fig. 2C) or the response to anti-PD-L1 immunotherapy (Fig. 2F) in patients with urothelial cancer.

Together, these results emphasize the importance of ITGAM and ITGAX expression levels in urothelial cancer progression and response to anti-PD-L1 immunotherapy.

### β2-integrin regulator expression levels effect on survival and response to immunotherapy

We further examined how the expression level of two-well known β2-integrin regulators, FLNA and TLN1, affected the overall survival and response to anti-PD-L1 immunotherapy in urothelial cancer.

Remarkably, patients with low levels of FLNA (FLNA low) had a significantly longer overall survival compared to those with high levels of FLNA (FLNA high) and reached median survival considerably later (almost 5-month delay) (Fig. 3A). They also responded significantly better to anti-PD-L1 immunotherapy (Fig. 3C). In contrast, different expression levels of TLN1 did not seem to significantly alter the overall survival of the patients (Fig. 3B) or their response to immunotherapy (Fig. 3D).

**Fig. 3.**
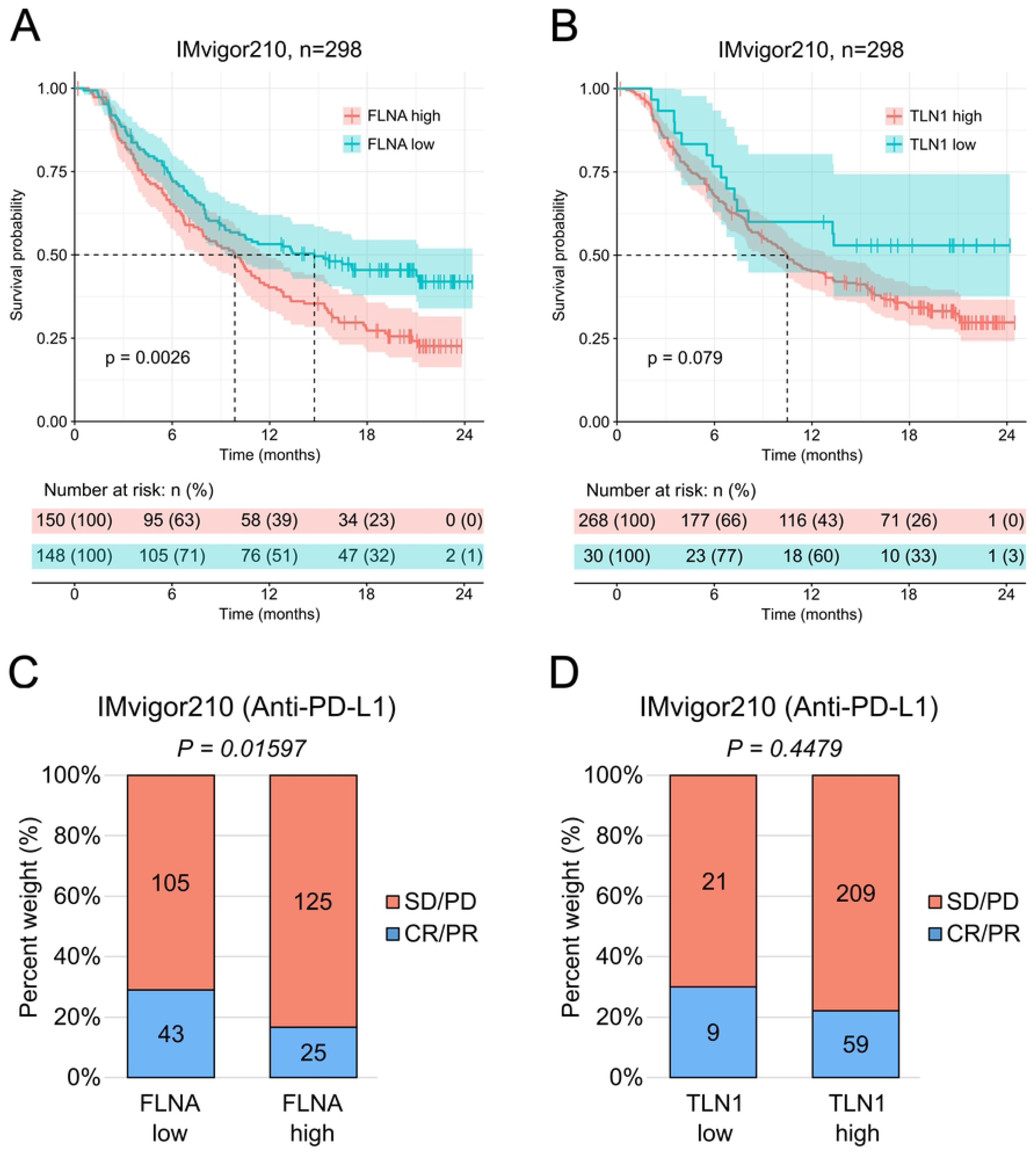
Low expression levels of FLNA correlate with longer overall survival in urothelial cancer patients and enhanced response to anti-PD-L1 immunotherapy. **A-B**, effect of gene expression in infiltrating leukocytes. Kaplan-Meier estimator of the overall survival after treatment with anti-PD-L1 of FLNA high (n=150) and FLNA low (n=148) patients (**A**), and TLN1 high (n=268) and TLNA low (n=30) patients (**B**). The dotted lines indicate the time at which each group reached median survival (FLNA high: 9.86 months, FLNA low: 14.75 months; TLN1 high: 10.48 months, TLN1 low: not applicable). The colored vertical marks on the plot indicate censored events for each group in time. Below the plot, risk table indicating the number of patients at risk belonging to each group at each point in time: high, in orange, or low, in blue. Statistical analysis was done using the log-rank (Mantel-Cox) test. P-values are shown on the plots. **C-D**, rate of clinical response to anti-PD-L1 immunotherapy (SD/PD, stable disease/progressive disease, n=150 (**C**), n=268 (**D**); CR/PR, complete response/partial response, n=148 (**C**), n=30 (**D**)). Statistical analysis was done using the chi-squared test. P-values are shown on the plots.

Together, these results underline a key role for FLNA expression levels in urothelial cancer progression and response to anti-PD-L1 immunotherapy.

### Association of ITGAL and FERMT3 expression levels with adaptive immune response genes

Our results indicated that different β2-integrins and their regulators have very different roles in urothelial cancer development. Given β2-integrins are immune-specific proteins, we sought to investigate their role in the immune landscape of the tumors. To this end, we performed differential gene expression analysis on each of the eight genes of interest, comparing the groups with low versus high expression levels.

We observed a significant correlation of β2-integrins and immune genes in the samples (Fig. 4). For example, downregulated markers in the ITGAL (Fig. 4B) and FERMT3 (Fig. 4D) low group revealed pathways such as “leukocyte activation”, “adaptive immune response”, “positive regulation of immune response”, “leukocyte migration”, “humoral immune response” and “regulation of cell killing”. For instance, genes such as CD2 (a costimulatory molecule involved in immunological synapse formation), ZAP70 (a tyrosine kinase which has an essential role in T cell activation), IL2RB (the IL-2 receptor which is essential for T cell function), BTLA (a lymphocyte co-signaling molecule of the CD28 superfamily), and KLRG1 (an inhibitory lectin-like receptor expressed on T cells and NK cells) were downregulated in ITGAL low group (Fig. 4A). Low expression of ITGAL and FERMT3 therefore appears associated with suppressed adaptive immune responses in the tumors.

**Fig. 4.**
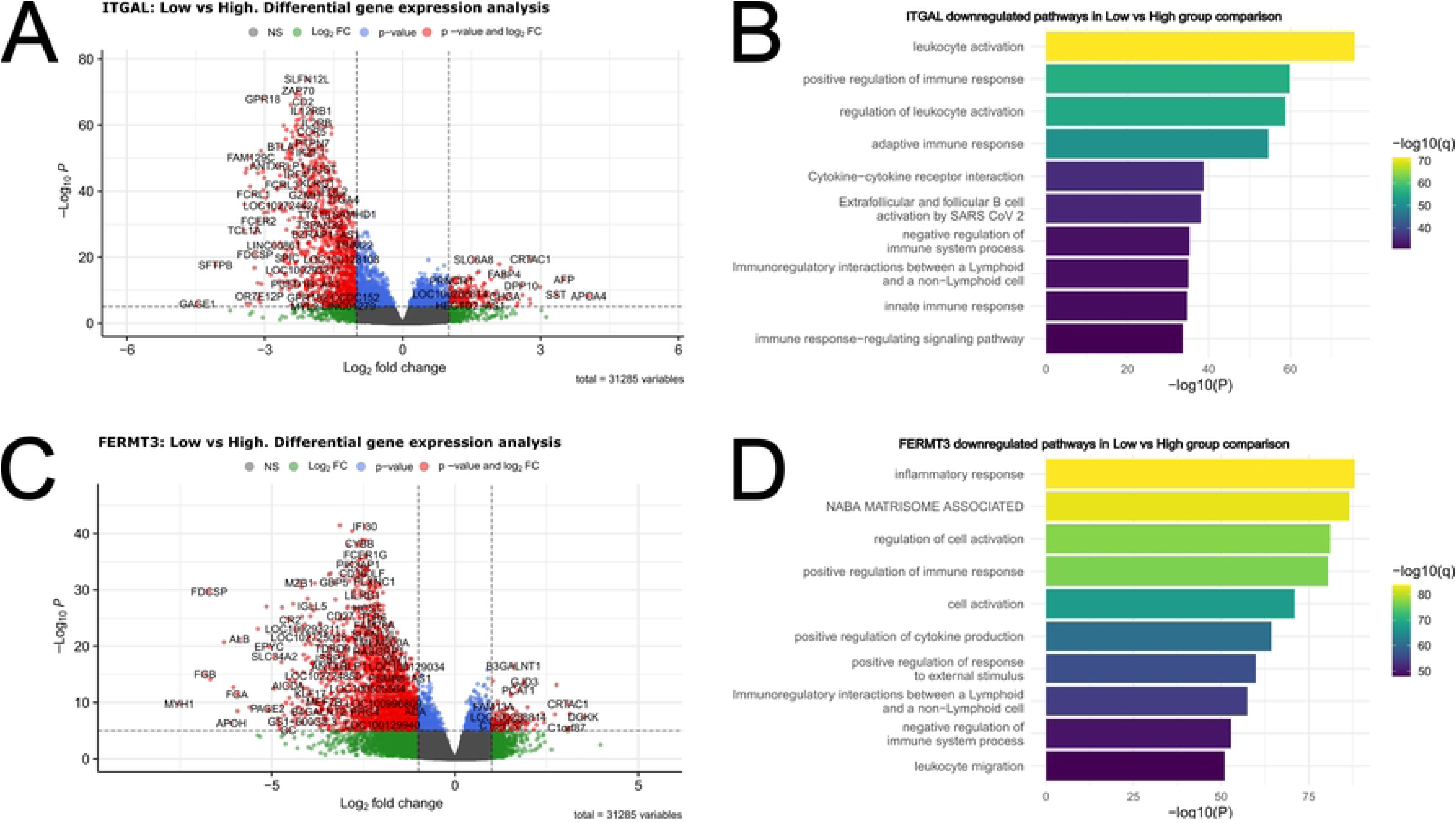
Differential gene expression analysis and gene set enrichment analysis for ITGAL and FERMT3. Volcano plots showing the differentially expressed genes in the group comparison low vs high for ITGAL (A) and FERMT3 (C). All genes that passed the p-value and log2 fold change threshold are indicated in red. Downregulated genes in the comparison have a negative Log2 fold change and upregulated genes have a positive Log2 fold change. Functionally enriched pathways were plotted for ITGAL (B) and FERMT3 (D).

### Association of ITGAM and ITGAX expression levels with myeloid immune response genes

In contrast to ITGAL and FERMT3, ITGAM and ITGAX showed negative association with survival, and also negative correlation with response to immunotherapy (Fig 2). The differential gene expression analysis again revealed association of these markers with immune genes (Fig. 5B and 5D), but the pathways identified were in part different from ITGAL/FERMT3. For instance, in the ITGAM low group, pathways such as “inflammatory response”, “neutrophil degranulation”, “myeloid leukocyte activation”, “innate immune response” and “phagosome” were downregulated compared to the ITGAM high group (Fig 5B). For example, OSCAR (a FcRγ associated receptor expressed in myeloid cells, involved in antigen presentation), CCL18 (an immunosuppressive, pro-tumorigenic chemokine produced by dendritic cells and macrophages, associated with M2 macrophage polarization), CHIT1 (an enzyme with important roles in macrophage biology), LTF and MPO (enzymes found in myeloid cells such as neutrophils) were downregulated in the ITGAM/ITGAX low groups (Fig. 5A and 5C). This analysis clearly revealed an association of ITGAM (and in part ITGAX) with the myeloid immune landscape of the tumors.

**Fig. 5.**
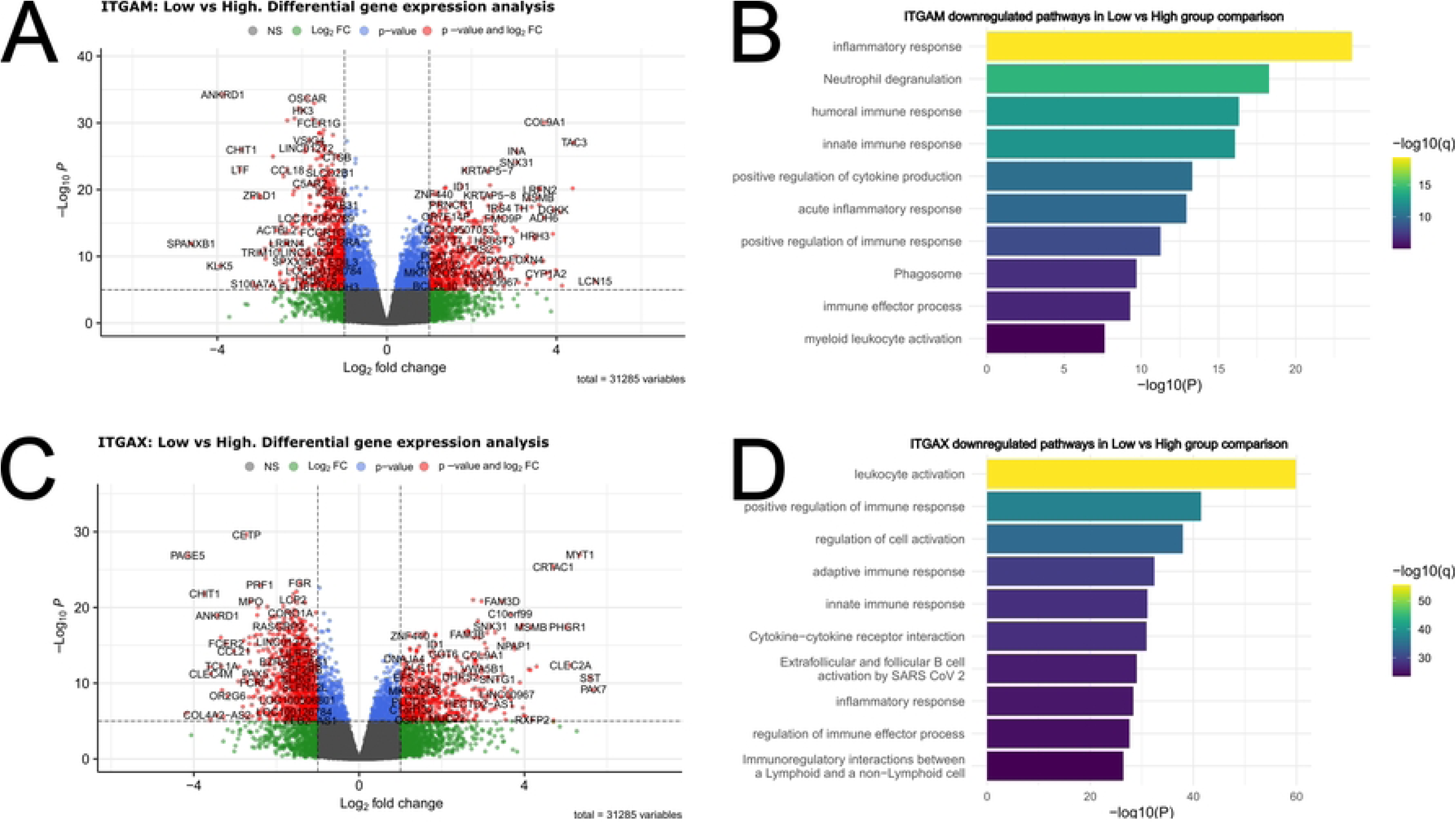
Differential gene expression analysis and gene set enrichment analysis for ITGAL and FERMT3. Volcano plots showing the differentially expressed genes in the group comparison low vs high for ITGAM (A) and ITGAX (C). All genes that passed the p-value and log2 fold change threshold are indicated in red. Downregulated genes in the comparison have a negative Log2 fold change and upregulated genes have a positive Log2 fold change. Functionally enriched pathways were plotted for ITGAM (B) and ITGAX (D).

### Association of FLNA expression levels with ECM genes

Filamin A is expressed in immune cells but also in other types of cells. Indeed, the differential gene expression analysis of FLNA did not reveal an association with immune genes; instead, in FLNA low samples pathways such as “NABA core matrisome”, “extracellular matrix organization”, “response to wounding” and “regulation of cell-substrate adhesion” were affected (Fig 6B). For example, MYL9 (myosin light chain 9, involved in focal adhesions and integrin-mediated cell adhesion/migration), ITGA5 (integrin α5), FLNC (filamin C, involved in Cell-extracellular matrix interactions and cell-cell communication), TGFB1I1 (Transforming growth factor β 1 induced transcript 1, involved in cell proliferation and migration) were downregulated in FLNA low samples (Fig. 6A). Thus, filamin A expression levels seem to regulate mainly the tumor extracellular microenvironment and not directly affect immune genes in the samples.

**Fig. 6.**
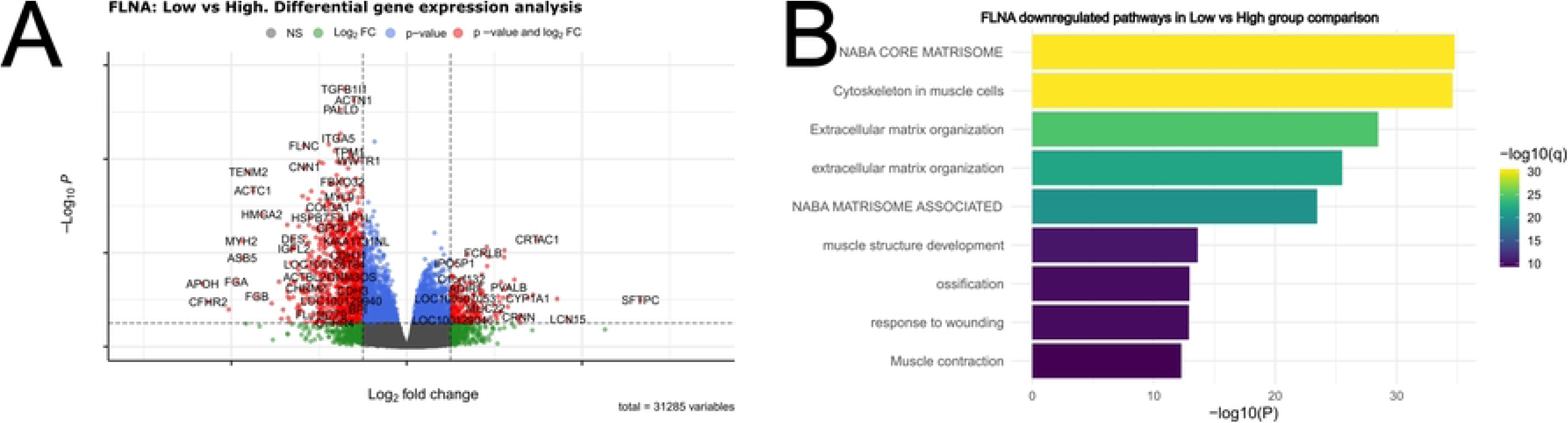
Differential gene expression analysis and gene set enrichment analysis for FLNA. Volcano plots showing the differentially expressed genes in the group comparison low vs high for FLNA (A). All genes that passed the p-value and log2 fold change threshold are indicated in red. Downregulated genes in the comparison have a negative Log2 fold change and upregulated genes have a positive Log2 fold change. Functionally enriched pathways were plotted for FLNA (B).

### Effect of **β**2-integrins and their regulators on the immune landscape in urothelial cancer

Our differential gene expression analysis indicated differences in the immune landscape in tumors expressing different levels of β2-integrins and their regulators. Next, we used EcoTyper(17) to see if stratified groups based on low and high expression of ITGAL, ITGAM, and FLNA correlated with differences in immune cell types within the tumor environment (Fig 7).

**Fig. 7.**
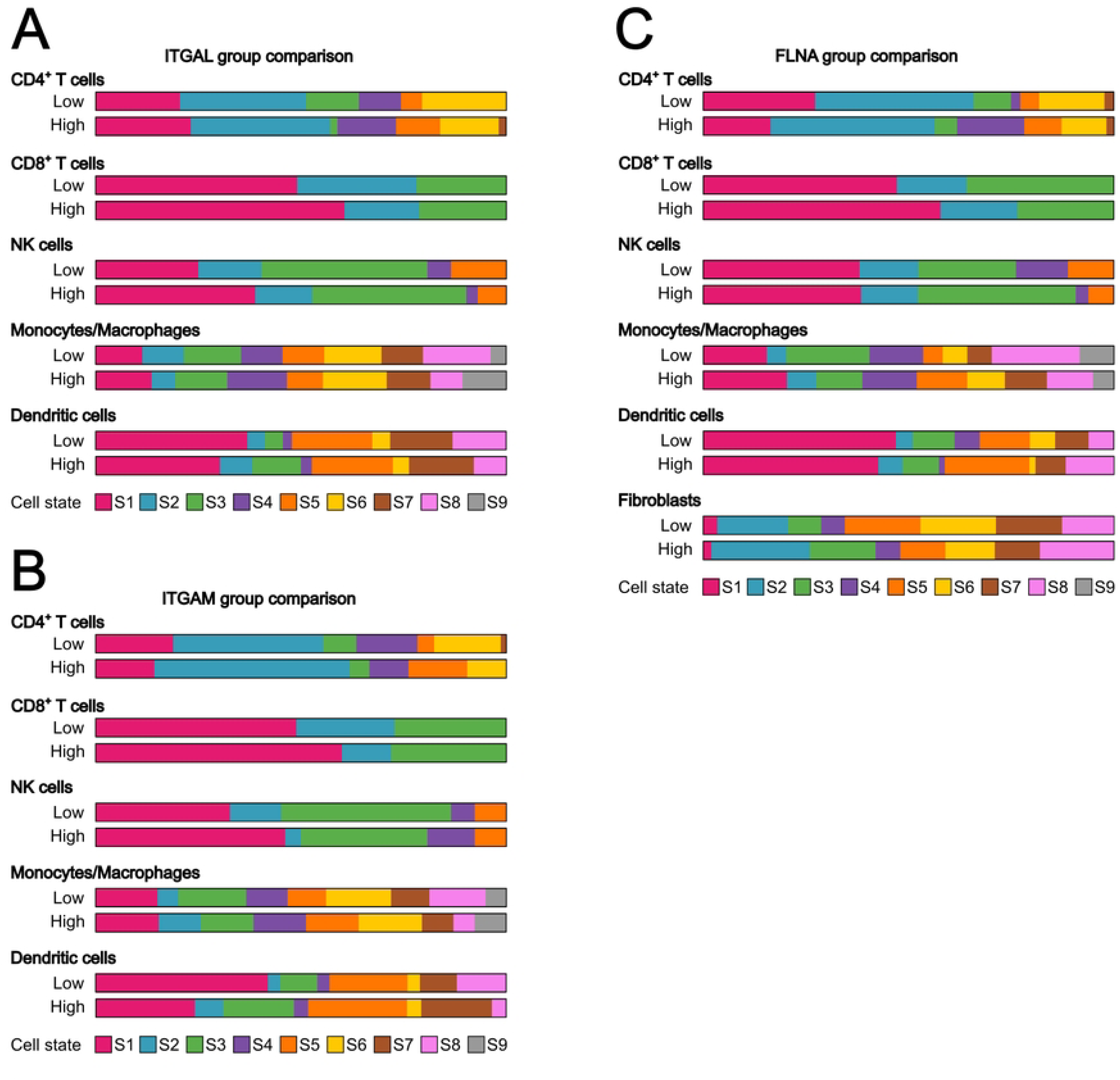
Comparison of cell states in ITGAL, ITGAM and FLNA groups. The proportion of different cell states of CD4+ T cells, CD8+ T cells, NK cells, Monocytes/Macrophages and Dendritic cells in the low expressing groups were compared to their corresponding high expressing groups for ITGAL (A), ITGAM (B), and FLNA (C), and are represented with colored bars. Cell states as defined by(17) are shown in the indicated colors.

Individuals belonging to the ITGAL low group (expressing a low amount of ITGAL, Fig 7A) showed lower amounts of Treg (S1, CXCR6^+^, CTLA4^+^) and naïve (CCR7^+^) CD4^+^ T cells, lower amount of naïve CD8^+^ T cells (S1, BTLA^+^, GZMK^+^) but higher amount of late-stage differentiated effector CD8^+^ T cells (S2, FCGR3A^+^, LAIR1^+^). Notably, the ITGAL low group of patients also had a lower amount of classical (PRF1^+^, CD247^+^) NK cells. As for myeloid cells, the ITGAL low group had a larger number of proliferative (S8, CDK4^+^), M0 (S2, FABP4^+^, MARCO^+^), M1 (CXCL9, SLAMF8^+^) macrophages, but smaller number of monocytes (S1, CCR2^+^, CLEC10A^+^), M2 (CD300E^+^, CLEC5A^+^) and M2-like (S1PR1^+^) macrophages. Dendritic cell amounts also differed, with the ITGAL low group having a higher amount of myeloid cDC1 (S1, CLEC9A^+^, XCR1^+^), but a lower amount of inflammatory myeloid cDC2-B (S2, ITGAM^+^, CLEC12A^+^) and mature immunogenic DCs (S3, PRF1^+^, CD274^+^, CD80^+^). ITGAL expression therefore significantly impacted on tumor infiltration of especially naïve CD8+ T cells and classical NK cells that have a positive correlation with survival (17).

When comparing the ITGAM groups (Fig 7B), we identified that the ITGAM low group had a higher amount of Treg (S1, CXCR6^+^, CTLA4^+^) and resting (KLF2^+^) CD4^+^ T cells. As for CD8^+^ T cells, they had a lower amount of naïve (S1, BTLA^+^, GZMK^+^), a bigger number of late-stage differentiated effector (S2, FCGR3A^+^, LAIR1^+^), but a similar number of effector memory (S3, IFNG^+^, GZMB^+^, LAG3^+^) CD8^+^ T cells compared to the ITGAM high group. NK cell pools were also different, with ITGAM low having less classical (S1, PRF1^+^, CD247^+^), but more normal-enriched (S2, STX11^+^) NK cells. Regarding myeloid cells, both groups had a similar number of monocytes (S1, CCR2^+^, CLEC10A^+^), but ITGAM low group had a lower amount of M0 (S2, FABPR4^+^, MARCO^+^), M2 (S4, CD300E, CLEC5A), and M2-like (S5, S1PR1^+^) macrophages than the ITGAM high group. ITGAM low had a larger number of M1 (S3, CXCL9^+^, SLAMF8^+^), and proliferative (S8, CDK4^+^) macrophages than the ITGAM high group. There were also differences in the types of dendritic cells, with ITGAM low group having a higher amount of myeloid cDC1 (S1, CLEC9A^+^, XCR1^+^) but lower amount of inflammatory myeloid cDC2-B (S2, ITGAM^+^, CLEC12A^+^), mature immunogenic (S3, PRF1^+^ CD247^+^, CD80^+^), mature (S5, CAV1^+^), and migratory activated (S7, CXCL2^+^, CXCL8^+^) dendritic cells. ITGAM expression therefore significantly affected especially the myeloid landscape of the tumors; In ITGAM low samples there were more M1 macrophages and myeloid cDC1 dendritic cells, that show a positive correlation with survival (17).

In the FLNA group comparison (Fig. 7C), the FLNA low group had more Treg (S1, CXCR6^+^, CTLA4^+^), slightly less naïve (S2, CCR7^+^), and substantially less resting (S4, KLF2^+^) CD4+ T cells than the FLNA high group. It also had less naïve (S1, BTLA^+^, GMZK^+^) and slightly less late-stage differentiated effector CD8^+^ T cells (S2, FCGR3A^+^, LAIR1^+^) than the FLNA high group. There were no big differences among the well characterized NK cell states. In the myeloid cell category, the FLNA low group had a lower number of monocytes (S1, CCR2^+^, CLEC10A^+^), M0 (S2, FABP4^+^, MARCO^+^), and M2-like (S5, S1PR1^+^), while it had a higher number of M1 (S3, CXCL9^+^, SLAMF8^+^), proliferative (S8, CDK4^+^) than the FLNA high group. Importantly, M2 foam cell-like macrophages (S6, AEBP1^+^), which have been recently shown to be significantly associated with patient survival (17), were decreased in this group. As for dendritic cells, the FLNA low group had slightly more myeloid cDC1 (S1, CLEC9A^+^, XCR1^+^), mature immunogenic (S3, PRF1^+^, CD274^+^, CD80^+^) and migratory-activated (S7, CXCL2^+^, CXCL8^+^), less mature (S5, CAV1^+^), and slightly less inflammatory myeloid cDC2-B (S2, ITGAM^+^, CLEC12A^+^) than the FLNA high group.

For FLNA we also analyzed fibroblasts; FLNA low group had especially less CAF2 (S2, CD34^+^, SPARCL1^+^), CAF1 (S3, POSTN^+^, COL10A1^+^) and pro-migratory-like (S8, CA9^+^) states, which are significantly associated with lower survival in cancer patients.

In conclusion, FLNA expression affects fibroblast phenotype, ECM gene expression and through that, appears to (indirectly) influence also especially myeloid immune landscape (M1/M2 macrophage balance, foamy cell macrophages) in the tumors.

### Correlation of **β**2-integrins and their regulators on carcinoma ecotype in urothelial cancer

We further used EcoTyper to analyze the carcinoma ecotypes(17) in the patient groups expressing different amounts of β2-integrins and their regulators (Fig 8). Carcinoma ecotypes (CEs) represent distinct transcriptional states of tumors that reflect underlying immune microenvironments and are strongly associated with clinical outcomes. These ecotypes serve as robust predictors of patient prognosis and response to immunotherapy, providing a powerful framework for stratifying tumors based on their immune and molecular characteristics. ITGAM high group tumors were enriched for CE1/CE2 carcinoma ecotypes, which are lymphocyte deficient and strongly linked to higher risk of death, in concordance with our findings that the ITGAM high group is associated with decreased survival. In addition, ITGAM high group tumors were also enriched for the CE3 ecotype, which is myeloid enriched and also associated with decreased survival (Fig. 8).

**Fig. 8.**
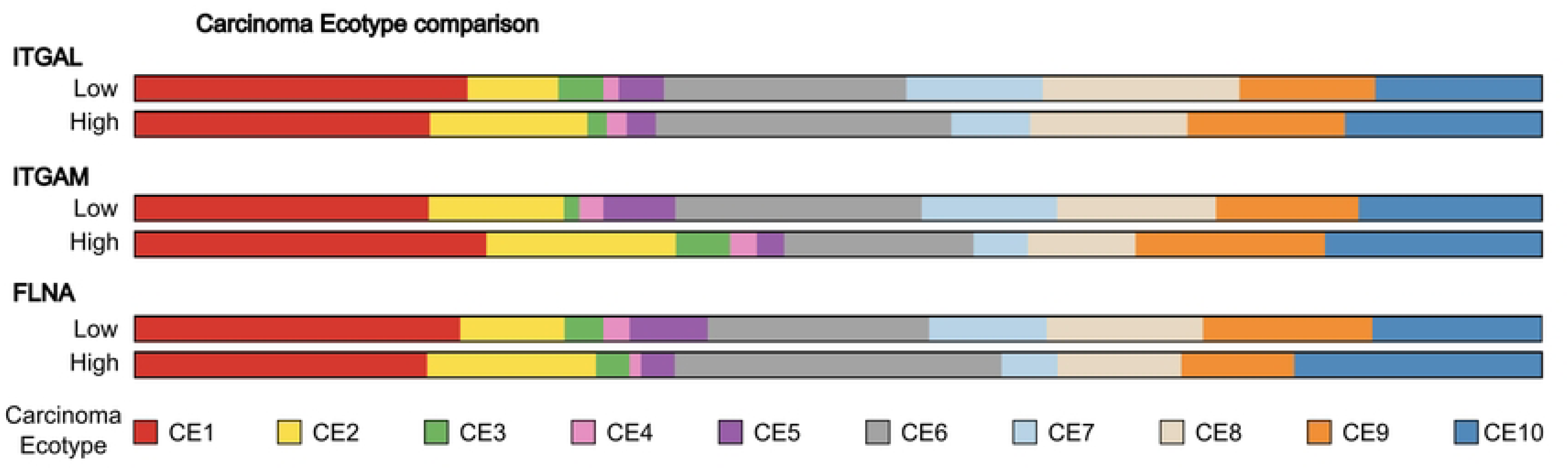
Carcinoma ecotypes of ITGAL, ITGAM and FLNA groups compared. The prevalence of the different carcinoma ecotypes in the low and high groups for ITGAL, ITGAM and FLNA was assessed and represented with colored bars. Carcinoma ecotypes as described by(17) re shown in the indicated colors.

The ITGAL high group tumors, which were associated with increased survival in the patients, showed reduced enrichment for the CE1 carcinoma ecotype and modest enrichment for immunogenic CE10 carcinoma ecotype, which is linked to naïve/central memory T cells, increased naïve B cell content and cDC1 dendritic cells. However, ITGAL high tumors did not show an enrichment of the CE9 ecotype (characterized by IFNγ signaling and T cell exhaustion), which is the best predictor of responsiveness to immunotherapy (Fig. 8). Interestingly, the FLNA low group, which did display an association with increased responsiveness to immunotherapy, did have an enrichment of CE9 carcinoma ecotype, showing that filamin expression, either directly or indirectly, does influence the immunological landscape of urothelial carcinoma (Fig. 8).

## Discussion

In this paper, we have conducted a series of bioinformatic analyses to investigate the role of the different β2-integrins in urothelial cancer and responsiveness to immunotherapy. This study reveals fundamentally different roles of different β2-integrins in tumor development and in determining the immune landscape of the tumors. ITGAL and FERMT3, encoding for the αL chain of LFA-1 and for kindlin-3, respectively, are positively associated with patient survival. Our gene expression and cancer Ecotyper analyses reveal that they appear to be most important in regulating adaptive (e.g. T cell mediated) responses to the tumors, but surprisingly, their expression level does not correlate with responsiveness to immunotherapy (although T cell infiltration is thought to be one of the most important determinants of anti-tumor immunity). A reason might be that ITGAL and FERMT3 expression may not correlate with T cell state, e.g. exhaustion, which is being targeted with immunotherapy approaches.

In contrast, myeloid genes e.g. ITGAM and ITGAX, show a negative correlation with survival, and regulate both the myeloid immune landscape of the tumors as well as their responsiveness to immunotherapy. Interestingly, FLNA, encoding for filamin A, an important regulator of integrins and cell adhesion, also regulated the immunological landscape of tumors, and its expression is inversely correlated with patient survival and response to immunotherapy. As filamin A is expressed both in immune cells and in other cell types, its effects on the immunological landscape may not be direct. Indeed, our gene expression and EcoTyper analyses suggest that filamin A expression affects fibroblast phenotype and the expression of extracellular matrix proteins, which may indirectly affect the immune profile of the tumors.

In conclusion, this study reveals that expression of specific β2-integrin subunits and regulators particularly ITGAL, FERMT3, ITGAM and FLNA may be used as biomarkers to differentiate urothelial cancer patients with different immune landscapes, survival profiles and responsiveness to immunotherapy.

## Methods

### Urothelial cancer data

Transcriptome-wide gene expression data and survival data was obtained from IMvigor210 urothelial cancer cohort, available via the R package IMvigor210CoreBiologies, license under the Creative Commons 3.0 and available for free download at http://research-pub.gene.com/IMvigor210CoreBiologies/. To ensure data completeness for survival analyses, 50 individuals lacking survival data information were excluded, resulting in 298 patients used for downstream analysis.

### Categorization of patients

To evaluate the association between gene expression levels and patient survival, the 298 patients from IMvigor210 cohort were categorized into “Low” and “High” expression groups for each gene of interest ITGB2, ITGAL, ITGAM, ITGAX, ITGAD, FERMT3, TLN1 and FLNA. This was done using the surv_cutpoint() function from the “survminer” R package (https://rpkgs.datanovia.com/survminer/index.html). The categorization employed log-rank statistics and accounted for patient survival, as done in(18–20). Thus, the continuous variable of gene expression for each of the genes of interest was transformed into a discrete variable and optimal cutoff values determined for each of the genes (Supplementary Table S1).

### Survival analysis

Survival analysis was done using the function survfit from the R package “survival” (https://cran.r-project.org/package=survival). Kaplan-Meier estimates of survival curves were plotted using the R package “survminer”. Statistical significance between “Low” and “High” expression level groups of each of the eight proteins of interest was determined using the log-rank (Mantel-Cox) test.

### Clinical response rate to anti-PD-L1 immunotherapy

Following the categorization of patients into “Low” and “High” expression level groups for each of the eight genes of interest, groups were compared based on their clinical response to immunotherapy (SD/PD, stable disease/progressive disease, CR/PR, complete response/partial response) and plotted. Statistical analysis was done using the chi-squared test.

### Differential gene expression analysis

Differential gene expression analysis was performed using “DESeq2” R package (http://www.bioconductor.org/packages/release/bioc/html/DESeq2.html). The differentially expressed genes (DEGs) were identified by comparing “Low” group vs “High” group at adjusted p-values < 0.05. The results were visualized using volcano plots using EnhancedVolcano R package (https://github.com/kevinblighe/EnhancedVolcano).

### Functional enrichment analysis

Functional enrichment analysis was performed to identify key biological processes and pathways linked to differentially expressed genes (DEGs) between the Low vs High conditions. Gene Ontology (GO) and Kyoto Encyclopedia of Genes and Genomes (KEGG) pathway analyses were mainly carried out using Metascape(21), which uses consolidated multiple up-to-date biological databases. DEGs were filtered based on an absolute log2 fold change threshold of |1.5| and a false discovery rate (FDR)-adjusted p-value ≤ 0.05, and only those meeting these criteria were included in the enrichment analysis.

### EcoTyper analysis

Gene expression data of the urothelial cohort standardized as transcript per million (TPM), and categorized into groups (low/high) for ITGAL, ITGAM, and FLNA was inputted to EcoTyper (https://ecotyper.stanford.edu/).

## Acknowledgements

We would like to thank Imrul Faisal and Daniel Davies for insightful discussions. This study was funded by Research council of Finland, Magnus Ehrnrooth foundation and Liv och Hälsa foundation (all to S.C.F.).

## Author contributions

S.C.F. and M.LL.A. designed the study. M.LL.A. performed analysis and wrote the paper together with S.C.F. I.K. performed analysis for the paper. All authors contributed to reviewing and editing of the manuscript.

## Data availability statement

Transcriptome-wide gene expression data and survival data was obtained from IMvigor210 urothelial cancer cohort, available via the R package IMvigor210CoreBiologies, license under the Creative Commons 3.0 and available for free download at http://research-pub.gene.com/IMvigor210CoreBiologies/.

## References

1. Sung H, Ferlay J, Siegel RL, Laversanne M, Soerjomataram I, Jemal A, et al. Global Cancer Statistics 2020: GLOBOCAN Estimates of Incidence and Mortality Worldwide for 36 Cancers in 185 Countries. CA Cancer J Clin. 2021 May 4;71(3):209–49.

2. Zhang T, Harrison MR, O’Donnell PH, Alva AS, Hahn NM, Appleman LJ, et al. A randomized phase 2 trial of pembrolizumab versus pembrolizumab and acalabrutinib in patients with platinum-resistant metastatic urothelial cancer. Cancer. 2020 Oct 15;126(20):4485–97.

3. Suzman DL, Agrawal S, Ning Y min, Maher VE, Fernandes LL, Karuri S, et al. FDA Approval Summary: Atezolizumab or Pembrolizumab for the Treatment of Patients with Advanced Urothelial Carcinoma Ineligible for Cisplatin-Containing Chemotherapy. Oncologist. 2019 Apr 1;24(4):563–9.

4. Fagerholm SC, Guenther C, Llort Asens M, Savinko T, Uotila LM. Beta2-Integrins and Interacting Proteins in Leukocyte Trafficking, Immune Suppression, and Immunodeficiency Disease. Front Immunol [Internet]. 2019/03/07. 2019;10:254. Available from: https://www.ncbi.nlm.nih.gov/pubmed/30837997

5. Harjunpaa H, Llort Asens M, Guenther C, Fagerholm SC. Cell Adhesion Molecules and Their Roles and Regulation in the Immune and Tumor Microenvironment. Front Immunol [Internet]. 2019/06/25. 2019;10:1078. Available from: https://www.ncbi.nlm.nih.gov/pubmed/31231358

6. Calderwood DA, Zent R, Grant R, Rees DJ, Hynes RO, Ginsberg MH. The Talin head domain binds to integrin beta subunit cytoplasmic tails and regulates integrin activation. J Biol Chem [Internet]. 1999/09/25. 1999;274(40):28071–4. Available from: https://www.ncbi.nlm.nih.gov/pubmed/10497155

7. Klapholz B, Brown NH. Talin - the master of integrin adhesions. J Cell Sci [Internet]. 2017/07/14. 2017;130(15):2435–46. Available from: https://www.ncbi.nlm.nih.gov/pubmed/28701514

8. Morrison VL, MacPherson M, Savinko T, Lek HS, Prescott A, Fagerholm SC. The beta2 integrin-kindlin-3 interaction is essential for T-cell homing but dispensable for T-cell activation in vivo. Blood [Internet]. 2013/07/05. 2013;122(8):1428–36. Available from: https://www.ncbi.nlm.nih.gov/pubmed/23823319

9. Moretti FA, Moser M, Lyck R, Abadier M, Ruppert R, Engelhardt B, et al. Kindlin-3 regulates integrin activation and adhesion reinforcement of effector T cells. Proc Natl Acad Sci U S A [Internet]. 2013/10/04. 2013;110(42):17005–10. Available from: https://www.ncbi.nlm.nih.gov/pubmed/24089451

10. Moser M, Bauer M, Schmid S, Ruppert R, Schmidt S, Sixt M, et al. Kindlin-3 is required for beta2 integrin-mediated leukocyte adhesion to endothelial cells. Nat Med [Internet]. 2009/02/24. 2009;15(3):300–5. Available from: https://www.ncbi.nlm.nih.gov/pubmed/19234461

11. Kiema T, Lad Y, Jiang P, Oxley CL, Baldassarre M, Wegener KL, et al. The molecular basis of filamin binding to integrins and competition with talin. Mol Cell [Internet]. 2006/02/04. 2006;21(3):337–47. Available from: https://www.ncbi.nlm.nih.gov/pubmed/16455489

12. Savinko T, Guenther C, Uotila LM, Llort Asens M, Yao S, Tojkander S, et al. Filamin A Is Required for Optimal T Cell Integrin-Mediated Force Transmission, Flow Adhesion, and T Cell Trafficking. J Immunol [Internet]. 2018/03/28. 2018;200(9):3109–16. Available from: https://www.ncbi.nlm.nih.gov/pubmed/29581355

13. Han C, Jin J, Xu S, Liu H, Li N, Cao X. Integrin CD11b negatively regulates TLR-triggered inflammatory responses by activating Syk and promoting degradation of MyD88 and TRIF via Cbl-b. Nat Immunol [Internet]. 2010/07/20. 2010;11(8):734–42. Available from: https://www.ncbi.nlm.nih.gov/pubmed/20639876

14. Varga G, Balkow S, Wild MK, Stadtbaeumer A, Krummen M, Rothoeft T, et al. Active MAC-1 (CD11b/CD18) on DCs inhibits full T-cell activation. Blood [Internet]. 2006/09/28. 2007;109(2):661–9. Available from: http://www.ncbi.nlm.nih.gov/pubmed/17003381

15. Guenther C, Faisal I, Fusciello M, Sokolova M, Harjunpaa H, Ilander M, et al. beta2-integrin adhesion regulates dendritic cell epigenetic and transcriptional landscapes to restrict dendritic cell maturation and tumor rejection. Cancer Immunol Res [Internet]. 2021/09/26. 2021; Available from: https://www.ncbi.nlm.nih.gov/pubmed/34561280

16. Morrison VL, James MJ, Grzes K, Cook P, Glass DG, Savinko T, et al. Loss of beta2-integrin-mediated cytoskeletal linkage reprograms dendritic cells to a mature migratory phenotype. Nature Communication. 2014;5.

17. Luca BA, Steen CB, Matusiak M, Azizi A, Varma S, Zhu C, et al. Atlas of clinically distinct cell states and ecosystems across human solid tumors. Cell. 2021 Oct;184(21):5482–5496.e28.

18. Schrock AB, Ouyang C, Sandhu J, Sokol E, Jin D, Ross JS, et al. Tumor mutational burden is predictive of response to immune checkpoint inhibitors in MSI-high metastatic colorectal cancer. Annals of Oncology. 2019 Jul;30(7):1096–103.

19. Fakih M, Ouyang C, Wang C, Tu TY, Gozo MC, Cho M, et al. Immune overdrive signature in colorectal tumor subset predicts poor clinical outcome. Journal of Clinical Investigation. 2019 Sep 16;129(10):4464–76.

20. Zhang Y, Xie R, Zhang H, Zheng Y, Lin C, Yang L, et al. Integrin β7 Inhibits Colorectal Cancer Pathogenesis via Maintaining Antitumor Immunity. Cancer Immunol Res. 2021 Aug 1;9(8):967–80.

21. Zhou Y, Zhou B, Pache L, Chang M, Khodabakhshi AH, Tanaseichuk O, et al. Metascape provides a biologist-oriented resource for the analysis of systems-level datasets. Nat Commun. 2019 Apr 3;10(1):1523.

